# The lack of myelin in the mutant *taiep* rat induces a differential immune response related to protection to the human parasite *Trichinella spiralis*

**DOI:** 10.1101/2020.04.02.021642

**Authors:** Karen Elizabeth Nava-Castro, Carmen Cortes, José Ramón Eguibar, Víctor Hugo Del Rio-Araiza, Romel Hernández-Bello, Jorge Morales-Montor

## Abstract

*Taiep* rat is a myelin mutant with a progressive motor syndrome characterized by tremor, ataxia, immobility episodes, epilepsy and paralysis of the hindlimbs, accompanied with differential expression of interleukins and their receptors that correlated with the progressive demyelination that characterize this mutant. Thus, the *taiep* rat is a suitable model to study neuroimmune alterations. The aim of this study was to investigate the immune alterations present in the mutant *taiep* rat during the acute infection with *Trichinella spiralis*. Our results show that there is an important decrease in the number of intestinal larvae in the *taiep* rat when compared to the Sprague-Dawley control rats. We also found differences in the percentage of innate and adaptive immune cell profile in the mesenteric lymphatic nodes and the spleen associated to the lack of myelin in the *taiep* rat. Finally, a clear pro-inflammatory cytokine pattern was seen in the infected *taiep* rat, which may explain the decrease in larvae number. These results sustain the theory that neuroimmune interaction is a fundamental process capable of modulating the immune response, particularly against the parasite *Trichinella spiralis* in a model of progressive demyelination that could be an important mechanism in autoimmune diseases and parasite infection.

**Author summary:** The complex communication among the brain and the immune system may be certainly altered during an infection and may be determinant in the resolution of this. We analyze the immune response to a parasite in a rat model in which a demyelinization process occur naturally and found that parasite loads were reduced when comparing with control subjects and this was accompanied to changes in the systemic immune response.

## Introduction

The interaction of the nervous, endocrine and immune systems is crucial in the maintenance of homeostasis in vertebrates and it is absolutely vital in mammals [1]. Moreover, hormones and neurotransmitters present in the immune cell microenvironment can restrict their autonomy, probably by acting on receptors of these neuroendocrine factors [2]. Therefore, efficient communication among these three systems implies the existence of afferent and efferent pathways, constituting a complex feedback system that regulates an adequate response in base of different insults.

Alterations of this network trigger pathologies that involve all the system components. In recent years, information on the multiple functions of the immune system has expanded remarkably; one of these functions has been biological adaptation through elimination of pathogens and foreign cells from the organism [3]. All functions require delicate control systems, which will allow adaptation of the organism to different physiological and pathological situations, which will face it a longer life. To meet this end, interaction with nervous and endocrine systems of the organism is necessary [3]. This interaction is constant and makes the merged functioning of the three systems. In other words, it involves common messengers and receptors, which simultaneously participate in a complex feedback system [4]. Alterations in the communication among the three systems lead to different pathologies. Numerous experimental data show cells of the immune system that are influenced by the neuroendocrine system, which displays various control levels, from metabolism to cell division, all of them regulated by hormones and neurotransmitters [5]. The immune response is possibly the only physiological phenomenon in which the amplification of the response is based on cell proliferation and specific transformation of their components [6]. This process requires metabolic changes and several growth factors, which makes the immune response dependent on neuroendocrine control [6].

*Taiep* rats were obtained as a spontaneous mutation during the inbreeding process to select the high-yawning (HY) subline of Sprague-Dawley rats [7]. This myelin mutant, developed a progressive motor syndrome characterized by: tremor, ataxia, immobility episodes, epilepsy and paralysis, the acronym of these symptoms given its name, *taiep* [7]. Importantly, tremor is present in both sexes when the mutants are 30 to 40 days old and it is characterized as a fine intentional tremor of the tail and proximal part of hindlimbs. By the fourth month, locomotor ataxia is quite evident and also changes in the walking pattern [8]. *Taiep* rats also showed a disorganized sleep-wake cycle along the circadian cycle. Therefore, during the light phase *taiep* rats had less total amount of REM sleep respect to normal Sprague-Dawley [9].

On the other hand, the *taiep* rat is an adequate model to analyze the neuroimmune interactions because the progressive demyelination of the Central Nervous System (CNS) is concurrent with immunological alterations. It is important to mention that astrocytes that surround blood vessels had abundant eNOS and because they are activated in *taiep* rats, it contributes to the higher levels of nitrates in the CNS on these myelin mutant animals [10, 11]. Additionally, there is also an infiltration of the microglia-macrophage lineage, which either express immunoreactive CD4 and CD8 positive cells being CD4-immunoreactive lymphocytes or activated microglia [10]. *Taiep* rats showed an increase in nitric oxide production, this characteristic is associated with lipoperoxidation and apoptosis in the cerebellum and brainstem, two brain regions that are the most demyelinated in this myelin mutant [12, 13].

Preliminary results showed that *taiep* rats have an increase in some cytokines, chemokines and their receptors in different structures of the CNS, such as brainstem, cerebellum and cerebral cortex that can participate in the survival of oligodendrocyte precursor cells (OPC) and improve re-myelination [14]. In fact, the levels of the chemokines CXCL1 (growth related oncogene, GRO) and CCL2 (monocyte chemoattractant protein-1, MCP-1) varied in their expression along the demyelination process in *taiep* rats and contribute to the deficit of myelination by OPC, similar to multiple sclerosis (MS) patients [14]. In the case of CCL5 (regulated on activation normal T cell expressed and secreted, RANTES) there is also a decreased in their expression that may contribute to a deficit in the communication between axon and glia, and could be responsible of demyelinated areas in the CNS of *taiep* rats [15]. As previously mentioned, all the immunological changes observed in *taiep* rats are in the CNS. However, so far, no description of changes of the immune system at systemic level, and much less the effect of the myelin deficiency on the outcome of an infection has been described.

*Trichinella spiralis* is an intracellular nematode that colonizes the striated muscles of infected mammals; this zoonosis, known as trichinosis, is commonly caused by the consumption of raw or undercooked meat from infected animals [16]. The establishment of the parasite in the duodenum represents the acute, or entherical, phase characterized by goblet cell hyperplasia, increased mucin and intestinal trefoil factor expression, and an inflammatory infiltration in the lamina propria [17]. At this stage, the intestinal inflammatory infiltrate is comprised of lymphocytes, mast cells, and eosinophils recruited to the intestinal Peyer patches and solitary lymphatic nodes [16]. Mastocitosis in the intestinal mucosa is also a typical feature of infection with *T. spiralis* [16], such activation of mast cells, followed by the secretion of its mediators, has been involved in the expulsion of parasites [17].

Despite the extensive research conducted on neuroimmunology, to date, no data has been published regarding whether progressive demyelination of the CNS affects duodenal integrity in response to gastrointestinal infection with *T. spiralis*. Thus, the aim of this study was to analyze the effects of *T. spiralis* infection in a naturally and progressive model of CNS demyelination, on the tissue components of the peripheral and local immune compartments. We also measure cell populations involved in immunity, as well as the cytokine levels produced and related with colonization during infection with *T. spiralis* and compare them with respect to Sprague-Dawley rats.

## Materials and Methods

### Ethics Statement

The protocol used in this study was approved by The Committee on Ethics and Use in Animal Experimentation of the Benemérita Universidad Autonóma of Puebla, and the Instituto de Investigaciones Biomédicas, UNAM. The study was done following the guidelines of Mexican regulations (NOM-062-ZOO-1999) and the Guide for the Care and Use of Laboratory Animals of the National Institute of Health, 8^th^ Edition to ensure compliance with the established international regulations and guidelines.

### Animals and experimental groups

Male Sprague-Dawley (SD) and *taiep* rats (around 300 gr) used in this study. The animals were single housed in a room with controlled temperature (22-24°C) and 12:12 light-dark conditions. The diet consisted of sterilized Harlan 2018 and sterile water *ad libitum*. The animals were organized into 4 groups of 5 animals each: 1) Intact control, 2) infected control, 3) *taiep* intact and 4) *taiep* Infected.

### Infection procedure

Infection procedure performed in base as previously described (Ibarra-Coronado et al, 2015). Briefly, consisted in inserting a gastric catheter, injecting 1,500 muscle larvae of *T. spiralis* suspended in 500μl of 1x PBS directly into the upper stomach. At the end of the experimental protocol, the rats were euthanized by anesthesia overdose (Sevorane®) and adult worms recovered by dissecting the small intestine into small sections, washed twice in 1X PBS, and incubated in sterile 1X PBS for 3 h at 37°C. Following incubation period, sedimented parasites were collected, washed in 1X PBS and quantified under a stereoscopic microscope (Velab Microscopes, Model VE-T2).

### Flow cytometry

Spleen and mesenteric lymph nodes (MLN) collected at the time of sacrifice were disaggregated using a sterile nylon mesh (70μm) and a syringe plunge in 1x PBS pH 7.4/4°C. The cell suspension was centrifuged at 182xg/3 min, decanted and resuspended in in 500 μl of erythrocyte lysis buffer (spleen only), incubated for 10 minutes at room temperature; 700 μl of FACS buffer were added followed by centrifugation at 182xg/3 min. Cells were resuspended in 500 μl of FACS buffer from which 25 μl were taken and 25 μl of primary antibody solution were added to the corresponding wells and incubated for 10 min/ 4°C. The following primary antibodies were used: AF-488 anti-rat CD3 (Biolegend. Clone 1F4, PE-Cy5 Mouse anti-rat CD4 (BD bioscience. Clone Ox-35), PE Mouse anti-rat CD8α (BD bioscience. Clone Ox-8), PE anti-rat CD45RA (Biolegend. Clone Ox-33), PE anti-rat TCRγδ (Biolegend. Clone V65) and AF647 anti-rat CD161 (Biolegend. Clone 1F4)). Wells were washed with 150 μl FACS buffer after incubation. Cells were finally resuspended in 100 μl of FACS buffer and 100 μl of fixation buffer (Paraformaldehyde 4% in PBS) and stored at 4°C in the dark. Data analysis was performed using the software FlowJo v7.6.

### Real time PCR for the determination of cytokines in spleen

Determination of the levels of mRNA expression of the genes of the cytokines pro-inflammatory (IL-1β, IL-6, TNF-α and IFN-γ,) and anti-inflammatory (IL-4, IL-5, IL-9, IL-13) were performed by PCR-quantitative in real time. Samples of splenic tissue were frozen in the TRIzol ®reagent (Ambion, ARN, Carlsband CA) at the time of slaughter. Total ribonucleic acid (RNA) was extracted with the same reagent, following the manufacturer’s protocol. First, the samples were transferred to sterile glass tubes containing 500 µL of TRIzol ® and the homogenized of the samples was made in the Polytron (all this was done at 4° C). Subsequently, the homogenization was transferred to a new tube that was added 200 µL of phenol-chloroform and 200 µL of water with Dietilpirocarbonate (H2O DEPC), stirred for 30 seconds and centrifuged at 13,000 rpm for 15 min at 4° C. The aqueous phase (containing the RNA) was recovered and transferred to a new tube. 200 µL of cold chloroform was added for each milliliter of aqueous phase recovered; the sample was stirred again and centrifuged at 13,000 rpm for 15 min a temperature 4° C. Again, the aqueous phase was recovered, and it was transferred to a new tube, where it was added cold isopropanol in a ratio1:1. It was stirred by investment and left at 4° C all night for the precipitation of the RNA to be carried out. The following day the samples were centrifuged at 13,000 rpm for 15 min at 4° C, decanted the supernatant being very careful not to peel the “pellet”, and this was washed with 1ml of ethanol (EtOH) at 75% cold. The sample was stirred until it was able to remove the pellet, and then it was centrifuged at 13,000 rpm for 15 min at 4° C. Again, the supernatant was decanted and centrifuged at 13,000 rpm for 1 minute at 4 ° C, in order to remove the remaining EtOH with a pipette.

Subsequently, the pellet was dried at room temperature until it took a translucent color and then resuspended in 30 µL of H2O DEPC, 7 ml were taken to make the quantification of RNA, and the gel of integrity. The RNA concentration was determined by absorbance at 260 nm, and integrity was verified after 1% agarose gel electrophoresis. The total RNA samples were immediately transcribed, using the MMLV-RT reverse transcriptase enzyme (Promega, Madison WI) to obtain the complementary DNA (cDNA) at a concentration of 5 mg/mL. To determine the levels of mRNA by PCR in real time, fluorogenic trials were performed using the reagent Light Cycler 480 SYBR Green I Master (Roche Applied Science, Mannheim, Germany). Fluorescence was detected in the equipment Light Cycler 480 Instrument (Roche Applied Science, Mannheim, Germany, Operator’s Manual Software Version 1.5).

Gene expression levels were determined by triplicate using 113 ng of the cDNA and the pair of oligos for each gene in plates 96 optical Wells LightCycler 480Multiwell plates 96/384 Clear (Roche Applied Science, Mannheim, Germany), using 18s and Cyclophilin as constituent controls. The sequence of the oligos sense and antisense that were used are presented in Table 1. The PCR amplification scheme used was 50 °c for 2 min, 95°c for 10 min, and 45 cycles of 94 °c for 15 seconds followed by 60 °c for 1 min. The relative expression was assessed in comparison with the reference Genes 18s and Cyclophilin, based on the following equation:

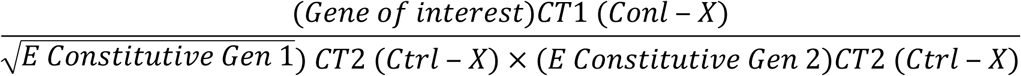

**Table 1.**
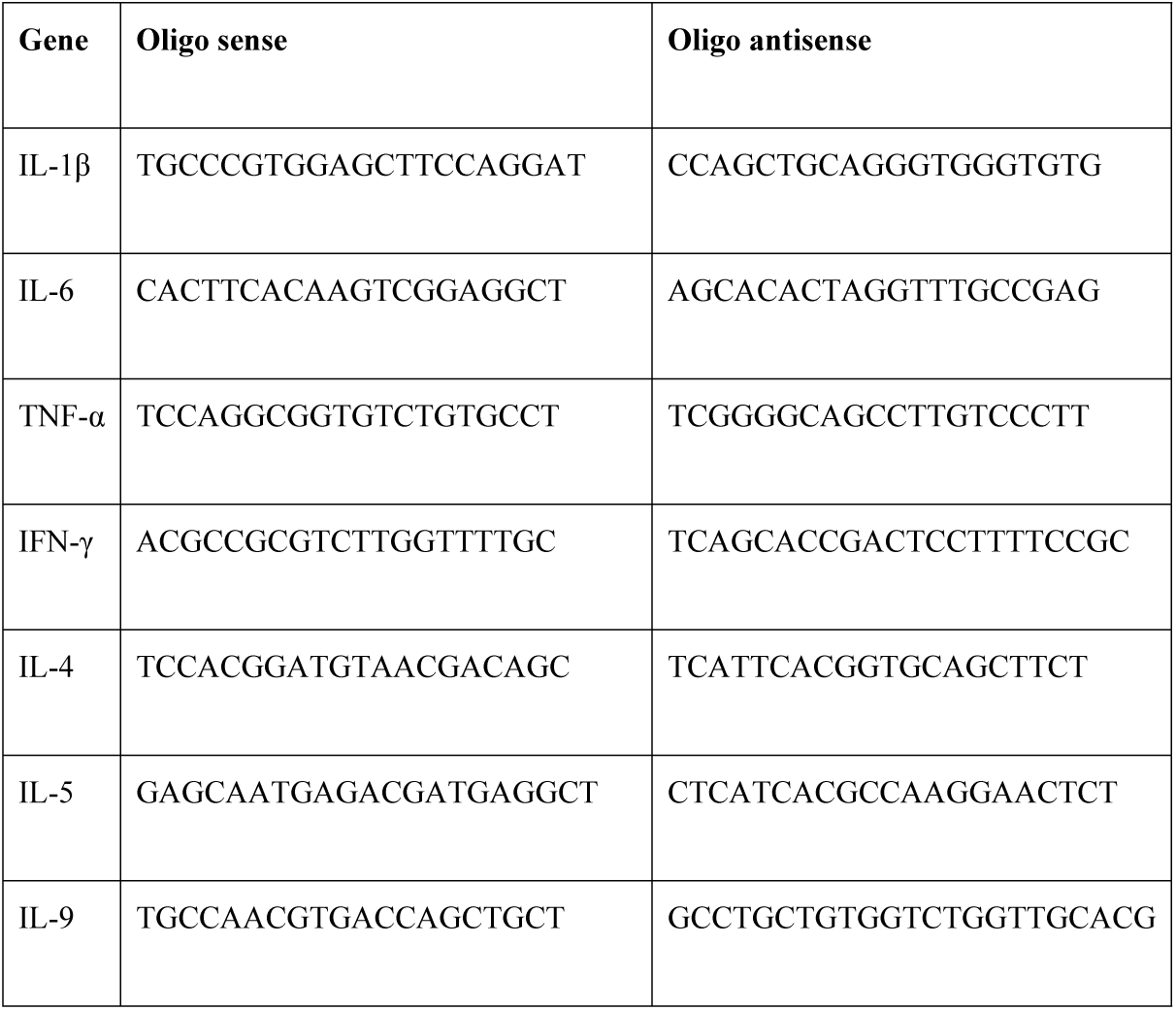
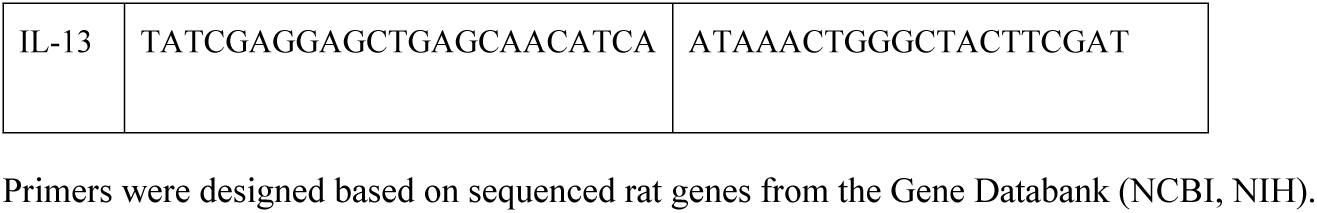
Oligonucleotide sequences used for RT-PCR.

### Statistical analysis and data processing

The general experimental design considers 2 independent variables: strain of rats (two levels: SD or *taiep*), and infection (Yes or No). Data from 2-3 independent experiments are charted as mean ± standard deviation and analyzed with Prism 6® software (GraphPad Software Inc.). Data distribution normality was assessed via Shapiro-Wilk test. Thereafter, a one-way ANOVA (α = 0.05) was performed, followed by a Tukey *post-hoc* test. Differences were considered significant when P < 0.05, with the actual probability P values being stated in each figure legend.

## Results

### Analysis of parasite burden

We observed that parasite burden is significantly decreased (3-fold) in the small intestine of the infected *taiep* rats, when compared with the outbred Sprague-Dawley (SD) infected rats (Figure 1); (p<0.001). Very few things, including anti-parasitic drugs, are reported to induce such a decrease in *T. spiralis* intestinal larvae number. This fact may suggest that lack of myelin may be related to intestinal function and immunity.

**Figure 1.**
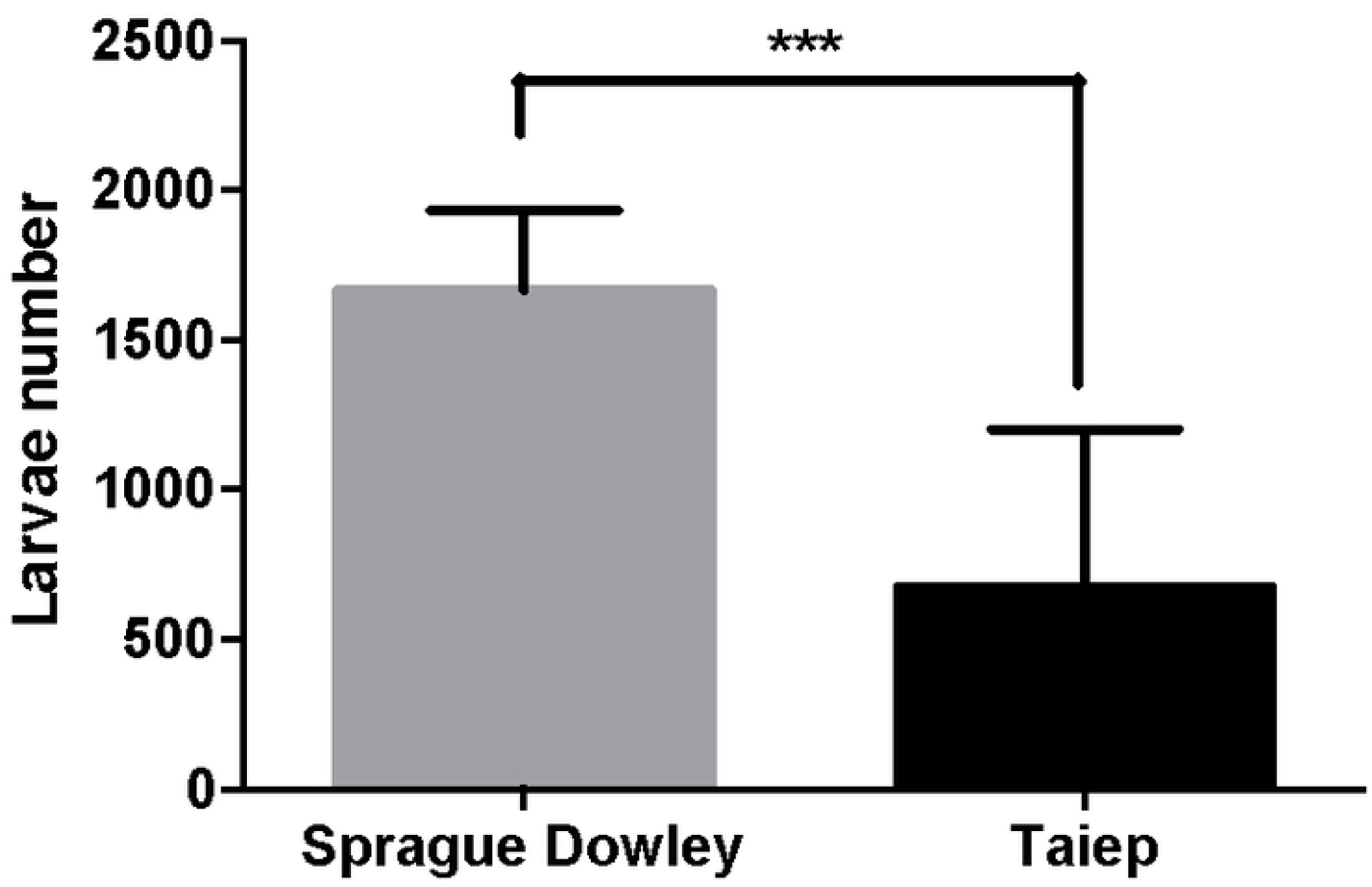
Parasite burden is significantly decreased in the small intestine of *taiep* infected rats. Animals were infected by *T. spiralis* larvae and 5 days after were sacrificed. In this figure, we represent outbred Sprague-Dawley rats as a control with respect to the mutant *taiep* rats. Data are the mean ± standard deviation of the mean. *** *P*<0.05

### Phenotypic characterization of immune cells in the spleen and mesenteric lymph nodes (MLN)

A phenotypic characterization of the innate and adaptive cells present in the spleen and mesenteric lymph nodes was performed. Figure 2 shows the gating pattern for the selection of the immune subpopulations. In panel A, lymphocytes were gated according to size and complexity and the expression of CD3 and CD45 markers were used to determine total T and B cells, respectively. In a second gating, CD3+ cells were gated and the expression of CD4 and CD8 markers were used to determine CD4^+^T (CD3^+^CD4^+^CD8^-^) T helper cells (Th) and CD8^+^T (CD3^+^CD4^-^CD8^+^) cytotoxic cells. Finally, in a third independent analysis, the expression of TCRγδ and CD161 were used to determine Tγδ cells and NK subpopulations. In panel B, macrophages selection was made by gating a bigger and more complex population (comparing to lymphocytes) and by the expression of CD11b.

**Figure 2.**
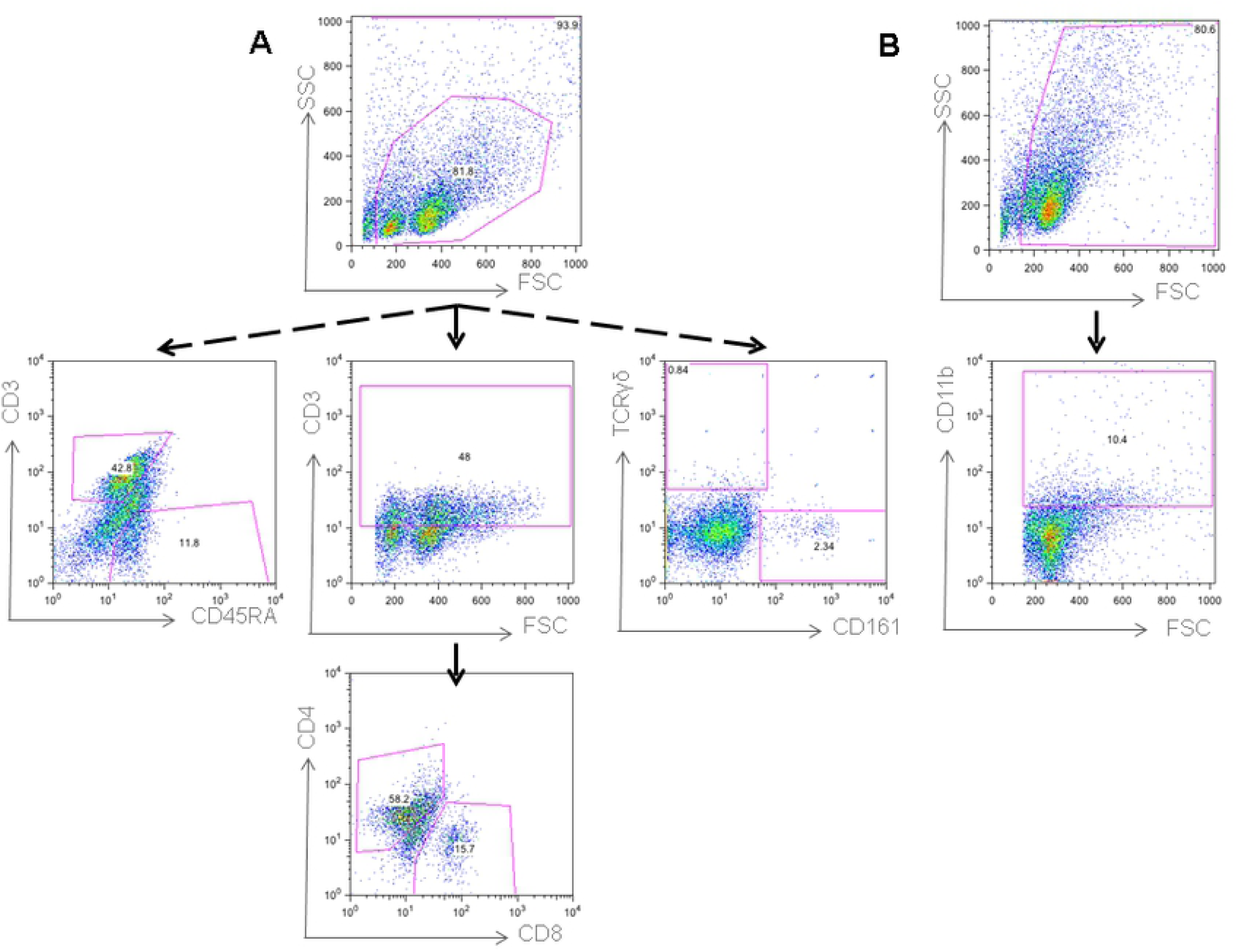
Gating pattern for the selection of the immune subpopulations in both spleen and mesenteric lymph nodes. (A) Lymphocytes were gated according to size and complexity. Total T and B cells and were gated by the expression of CD3 and CD45RA respectively. For T helper and T cytotoxic cells, a first gate was made in the CD3+ and then selected by the expression of CD4 and CD8 markers (CD3^+^CD4^+^CD8^-^) and (CD3^+^CD4^-^CD8^+^). Tγδ cells and NK cells were selected gated by the expression of TCRγδ and CD161. (B) Macrophages were selected by gating in a bigger and more complex population (comparing to lymphocytes) and by the expression of CD11b (Representative dot plots were used).

### Innate immune percent cell pattern in the MLN

When innate immune cells were analyzed in the MLN, we found no differences in the percentages of NK (Figure 3A) or Tγδ (Figure 3B). However, there was a significant increment in the percentage of macrophages in the *taiep* rats that was induced by the infection (p<0.05). These differences are not clear in the SD rats.

**Figure 3.**
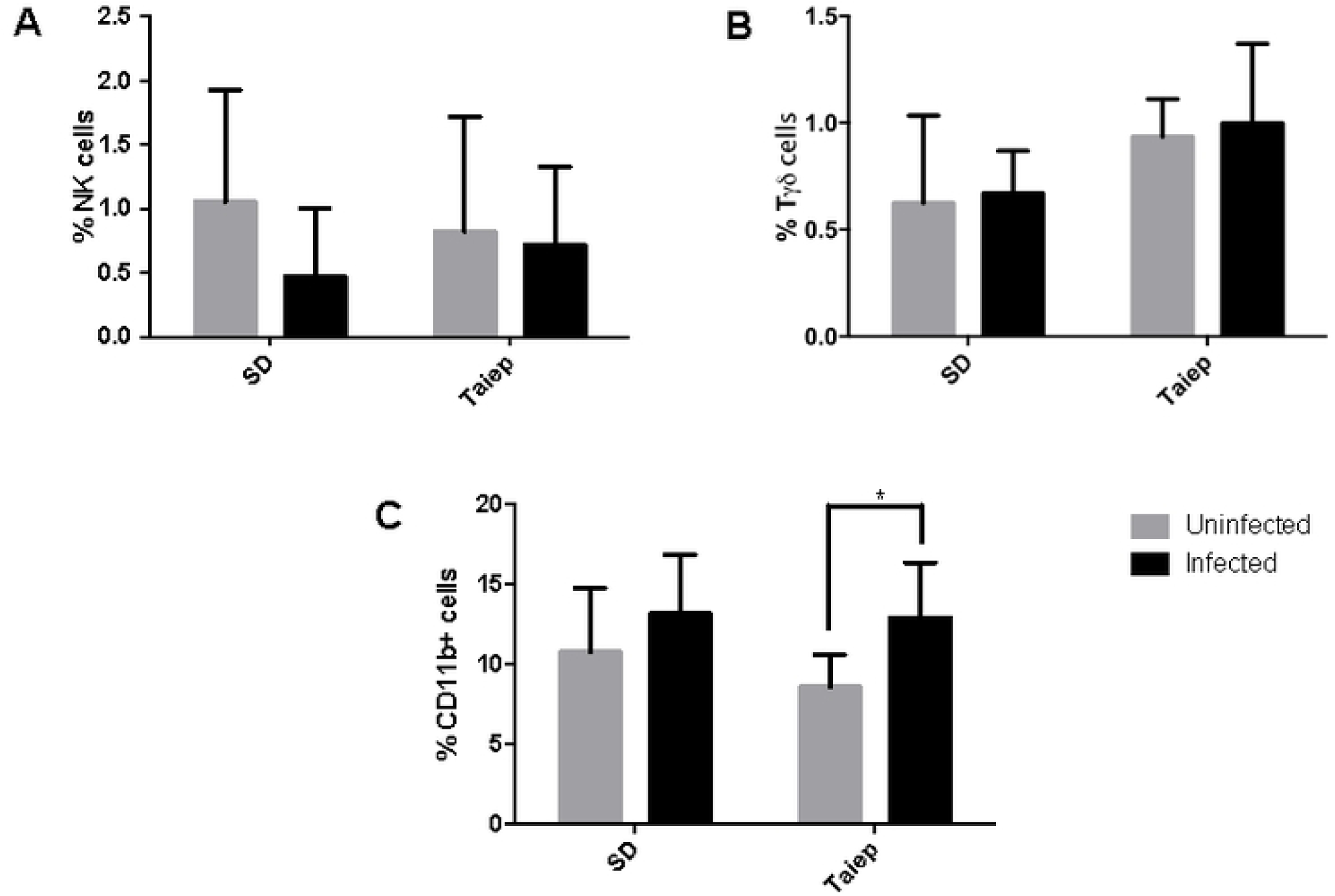
Analysis of the innate immune cells in mesenteric lymph nodes. Determination of immune subpopulations frequencies by flow cytometry in MLN of both SD and *taiep* rats with or without infection. Percentage of (A) NK cells (B) Tγδ and (C) Macrophages in non-infected (gray bars) and infected (black bars) Sprague-Dawley (SD) and *taiep* rats. Two-way ANOVA followed by Bonferroni test* *P*<0.05.

### Adaptive immune cell profile in the MLN

When T and B cells were analyzed in the MLN of SD and *taiep* rats, we found that infection induced a reduction in the percentage of total T cells in both SD and *taiep* rats (Figure 4A); and was observed equally for the CD4 T cell population (Figure 4C). In contrast, there was an increment in the percentage of B cells after infection. Interestingly, this increment caused by the infection was lower in the *taiep* rats that in the SD (p<0.01). There were no differences in the percentage of CD8 T cells.

**Figure 4.**
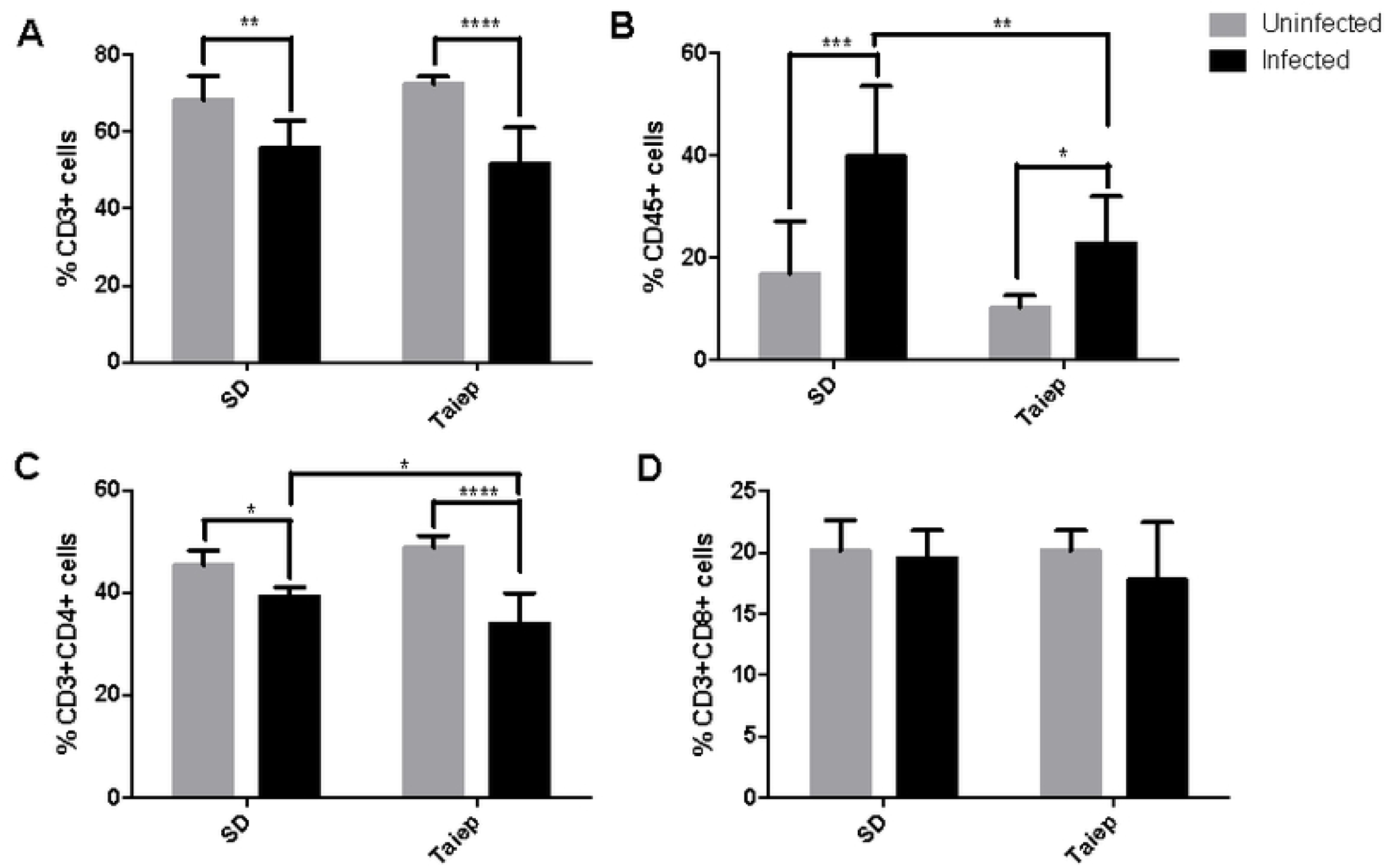
Analysis of the adaptive immune cells in mesenteric lymph nodes. Determination of immune subpopulations frequencies by flow cytometry in MLN of both SD and *taiep* rats with or without infection. Percentage of (A) Total T cells (B) B cells; (C) T helper cells and D) T cytotoxic cells in non-infected (gray bars) and infected (black bars) Sprague-Dawley (SD) and *taiep* rats. Two-way ANOVA followed by Bonferroni test \**p*<0.05; ** *p*<0.01, *** *p*<0.001.

### Characterization of the immune innate cell subpopulation profile in the spleen

We look for changes in the immune cell subpopulations in the spleen in order to determine changes in the central immunity. We look for differences in both innate and adaptative subpopulations as shown for the MLN. As expected, there were no differences in the percentages of NK (Figure 5A) or Tγδ (Figure 5B) in the spleen. As observed in the MLN, there was a slight increment in the resident macrophage subpopulation in the infected *taiep* rats when comparing with its non-infected counterparts. This increment was not significant in the SD rats (Figure 5C).

**Figure 5.**
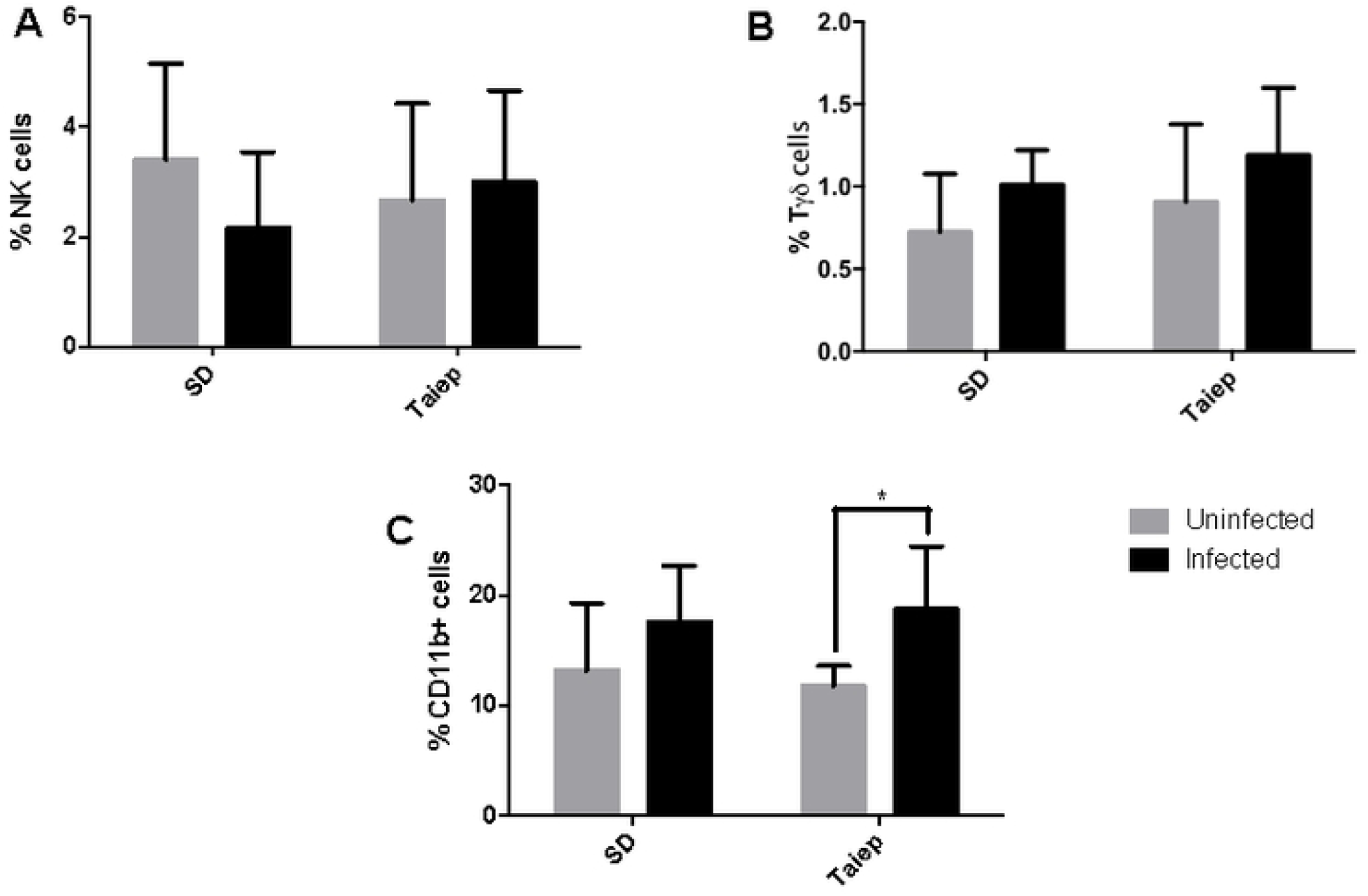
Analysis of the innate immune cells in the spleen. Determination of immune subpopulations frequencies by flow cytometry in MLN of both SD and *taiep* rats with or without infection. Percentage of (A) NK cells (B) Tδ and (C) Macrophages in non-infected (gray bars) and infected (black bars) Sprague-Dawley (SD) and *taiep* rats. Two way ANOVA followed by Bonferroni test * *P*<0.05.

### Adaptive immune cell profile in the spleen

In contrast with the reduction of the T cell population in the MLN, we found that the infection induced an increment in T cells in both SD and *taiep* rats. However, it is important to notice is that the percentage of the T cells in the *taiep* rats in lower quantities that the one observed in the SD (Figure 6A). When CD4 and CD8 T cells were analyzed, we found no differences caused by the infection and neither in SD nor *taiep* rats. In terms of the B cell population, we found that *taiep* rats showed a lower increment in the percentage of this population after infection when compared to SD rats (Figure 6B).

**Figure 6.**
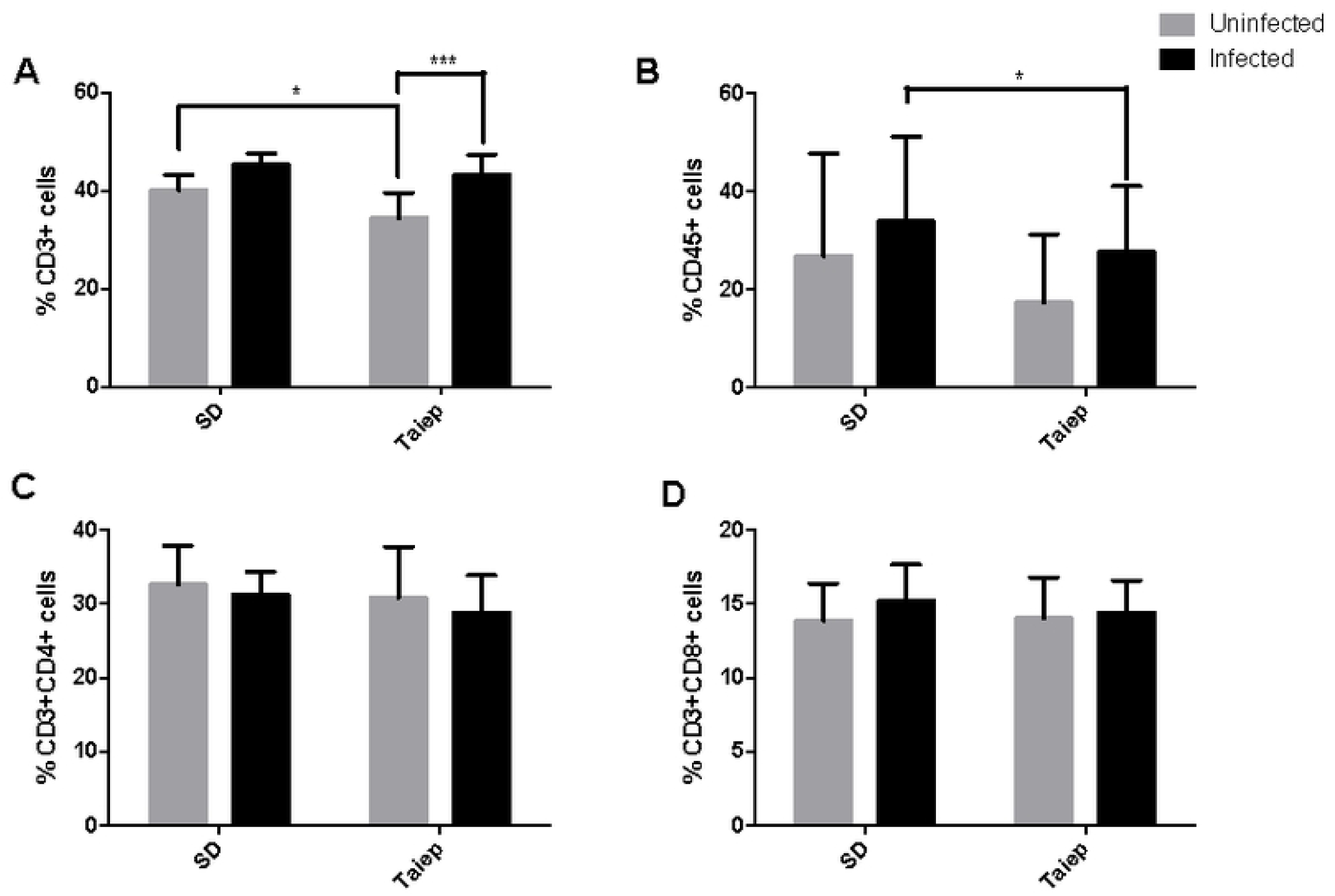
Analysis of the adaptive immune cells in the spleen. Determination of immune subpopulations frequencies by flow cytometry in MLN of both SD and *taiep* rats with or without infection. Percentage of (A) Total T cells (B) B cells; (C) T helper cells and D) T cytotoxic cells in non-infected (gray bars) and infected (black bars) Sprague-Dawley (SD) and *taiep* rats. Two-way ANOVA followed by Bonferroni test \**p*<0.05; *** *p*<0.001.

### Relative expression of cytokines Th1/Th2 in spleen

Since differences in the percentage of the different cellular subpopulations of the immune system does not necessarily reflect its function. The humoral mechanisms of these cells were continued to be evaluated by the determination of the expression of Th1 cytokines (IL-2 and IL-12), Th2 (IL-4 and IL-5), pro-inflammatory (IFN-γ and TNF-α and regulatory (IL-10 and TGF-β) at splenic level (Table 2). As for the production of cytokines Th1 can be observed that there are no changes in the expression of IL-2 and IL-12 (P > 0.05), (Table 2). However, the expression of IL-4 and IL-5 is increased due to infection in the *taiep* groups (p < 0.05), (Table 2). In the case of IFN-γ and TNF-α only an increase in its expression is observed due to infection in the *taiep* group (p < 0.01), (Table 2). In the case of both regulatory cytokines, IL-10 and TGF-β, they show a decrease that is significant (* p < 0.05) due to the effect of the infection. This decrease it is even more attenuated in the *taiep* rat (** p < 0.05) compared to infected rats.

**Table 2.** Cytokine pattern in spleen of uninfected (Intact) and infected Sprague-Dawley (Infected) and *taiep* uninfected and *taiep* infected rats.

|  |  |  | **Groups** |  |  |
| --- | --- | --- | --- | --- | --- |
| *Cytokine Family* | *Cytokine*<br>*Relative*<br>*Expression* | *Sprague-*<br>*Dawley* | *Infected* | *taiep*<br>*UNINFECTED* | *taiep*<br>*INFECTED* |
| Th1 | IL-2 | 9.3 ± 1.9 | 7.8 ± 2.1 | 6.1 ± 0.5 | 9.3 ± 0.9 |
|  | IL-12 | 5.0 ± 0.8 | 4.9 ± 1.1 | 5.2 ± 1.1 | 4.9 ± 1.1 |
| Th2 | IL-4 | 11.2 ± 2.8 | 18.89 ± 10.8 | 20.4 ± 2.5 ** | 59.89 ± 10.8 |
|  | IL-5 | 18.9 ± 3.1 | 29.0 ± 0.9 | 21.0 ± 2.8** | 70.0 ± 0.9 |
| Pro-inflammatory | IFN-γ | 6.3 ± 0.5 | 16.0 ± 1.1 | 18.6 ± 3.5** | 86.0 ± 1.1 |
| | TNF-α | 10.9 ± 1.0 | 21.0 ± 0.9 | 23.6 ± 3.3** | 100.0 ± 7.9 |
| Regulatory | IL-10 | 16.3 ± 0.4 | 20.1 ± 0.5 | 8.6 ± 1.6** | 13.9 ± 1.5 |
| | TGF-β | 26.3 ± 0.7 | 19.2 ± 1.9 | 7.2 ± 1.2** | 11.9 ± 1.1 |

Relative expression (Optical Density (OD) cytokine /OD control gene (18s). Data are presented as RE ± standard deviation. \*P<0.05, \*\*P<0.01.

## Discussion

The aim of the present study was to evaluate the effect of progressive demyelination in the *taiep rat*, on the systemic immune response and its association to the establishment of the gastrointestinal *parasite T. spiralis*. Although recent research on the effects of interrupted neuroimmune communication on physiological functions in experimental models, on metabolism [18], endocrine function [19] and interesting on immune functions [20, 21], has been carried out, so far this is the first study exploring the effect of natural myelin deprivation on the immune response against a gastrointestinal parasite. In this study, we addressed the influence of the demyelination of the CNS on the behavior of different cell populations at different levels of organization, which result in an alteration in the homeostasis of the immune system and therefore lead to changes in the mounted immune response. Our study provides evidence of how neuroimmune communication modifies the behavior of different cell populations at different levels of organization that impact the amount of infestation of a well-known parasite. Our results provide insight into the mechanisms by which neuroimmune interactions alter the acquisition of immunity against the human nematode parasite *T. spiralis* [22]. A study of the effects of myelin deprivation not only gives us knowledge of the role that myelin may have on different systems in the organism, but it also gives us tools to understand the relationship between different systems, in particular the CNS and immune system, which maintains a balance between them [23]. Understanding the interaction of these systems requires that we include all members of the participating systems. Thus, we cannot study the effect of neuroimmune interactions on an immune system that remains static, so we demonstrated the effect on physiologically or pathological relevant conditions, that is, reacting against an antigenic challenge and interacting with other systems, in our case the CNS deprived of myelin. Despite our extensive work studying the neuroendocrine system and the immune response, in the context of an antigenic challenge, we must also take into account the influence of infection on other systems [24, 25].

Furthermore, we have shown that alterations in the myelination of the CNS has a differential effect on the different components of the immune system; in particular, it can increase the concentration of NK cells and cytotoxic T cells, as well as pro-inflammatory mediators, such as IL-1β, TNF-α and IFN-γ. However, little is known about how myelin deprivation can alter the immune response generated in earlier periods of infection. Our findings show that demyelination of the CNS modifies immune cell percentage, and its cytokine pattern at systemic level, which impact on the number of larvae established in the duodenum. These changes may be able to modify certain behaviors to induce protection during acute infection and impair *T. spiralis* life cycle [22]. However, this field still remains poorly studied, and consequently the function of the CNS and cell components in the immune response to pathogens it is beginning to be understood. Talking about the CNS, a wide variety of research has addressed the importance of physiological functions, such as sleep for the functioning of the immune system [22, 26]. In 2009, Preston and colleagues analyzed the possibility that sleep has evolved across species to allow the organism special protection against parasitic infections [27]. These authors also analyzed the correlation between sleep duration and parasitic infection levels as an indicator of the number and type of parasites that are able to infect twelve mammalian species. In this case, a negative correlation was found, with a longer duration of sleep significantly correlating with a lower level of parasitic infection. Thus, authors conclude that sleep evolved to protect animals from parasitic infections [27]. Sleep is also altered in the *taiep* rats, because they had a disorganized sleep-wake cycle and during immobility episodes a rapid eye movement sleep emerges [9]. So probably sleep alterations it is also involved in the protection seen in this study against parasite load in the *taiep* rat. In this aspect our work addressed the effect of progressive myelin loss on the proportion of immune cells in the spleen and MLN at baseline (non-infection) and during a parasitic infection, in both Sprague-Dawley and *taiep* rats. To this end, the parasite *Trichinella spiralis* provided a suitable model of infection to evaluate the immune response that is generated in a short period of infection. The changes observed in the MLN give us the first evidence of the importance of strong changes in the CNS, such a diminution of myelin on cell recruitment into the peripheral lymph during the innate immune response. Furthermore, the analysis conducted in the spleen cell populations, presents an overview that suggests that hypomyelination and progressive demyelination in the CNS modifies the behavior of cells at the systemic level in basal conditions as well as during infection. In this respect, our results showed that in the spleen (reflects systemic effects), the infection status is irrelevant because the infection does not cause a differential effect on the changes observed, that could be due to the changes on myelin levels, so it is clear that the two stimuli produce different responses. Our correspondence analysis revealed that some cell populations respond similarly, but others do not share the same behavior. Overall, both TAIEP and SD rats are associated with an increase in the populations of B cells (CD45+) and NK cells (NK1.1+) in the spleen, and stress is also associated with increased total T cells (CD3+), but in contrast, the SD condition is associated with a decrease in the cytotoxic T cell subpopulation (CD8+CD3+). These results are in agreement with those from Velazquez-Moctezuma, 2004, which reported similar changes with different stressors, and therefore, we can postulate that the overall response of the cell subpopulations depends on the immune system nature of the stressor and the basal condition of the CNS and in this case myelin loss [28]. The major differences of *taiep* rats in comparison with Sprague-Dawley are that cytokines such as IFN-γ, TNF-α, and were unaltered but IL-1β and IL-10 were downregulated in 1-month-old *taiep* rats. The decrease in TGF-β1 expression at 6-month-old *taiep* and SD rats might also be a sign of CNS aging and demyelination in the former and only the first component in the second. It is probably that the alteration in microtubule proteins in the *taiep* rats causes protection against the infection with the human nematode *Trichinela spiralis*.

However, all the immune mechanisms involved in this protective effect are still unknown. Our results strongly suggest that the deficiency of the remyelination contributes to the increased immune function. Perhaps, another study may be designed to check on the specific site of the infection, the duodenum. Further studies are needed to identify signal pathways and other mediators involved in the immune response of the *taiep* rat, but our results clearly demonstrated that parasitic infection load depends on the neuroimmune interactions that took place in the demyelination process of *taiep* rats.

## Data Availability Statement

The raw data supporting the conclusions of this manuscript will be made available by the authors, without undue reservation, to any qualified researcher.

